# Dynamics of transcription-dependent H3K36me3 marking by the SETD2:IWS1:SPT6 ternary complex

**DOI:** 10.1101/636084

**Authors:** Katerina Cermakova, Eric A. Smith, Vaclav Veverka, H. Courtney Hodges

## Abstract

SETD2 contributes to gene expression by marking gene bodies with H3K36me3, which is thought to assist in the concentration of transcription machinery at the small portion of the coding genome. Despite extensive genome-wide data revealing the precise localization of H3K36me3 over gene bodies, the physical basis for the accumulation, maintenance, and sharp borders of H3K36me3 over these sites remains rudimentary. Here we propose a model of H3K36me3 marking based on stochastic transcription-dependent placement and transcription-independent spreading. Our analysis of the spatial distributions and dynamic features of these marks indicates that transcription-dependent placement dominates the establishment of H3K36me3 domains compared to transcription-independent spreading processes, and that turnover of H3K36me3 limits its capacity for epigenetic memory. By adding additional terms for asymmetric histone turnover occurring at transcription start sites, our model provides a remarkably accurate representation of H3K36me3 levels and dynamics over gene bodies. Furthermore, we validate our findings by revealing that loss of SPT6 impairs the transcription-coupled activity of the SETD2:IWS1:SPT6 ternary complex, thereby reducing the tight correlation between transcription and H3K36me3 levels at gene bodies.

## Introduction

Eukaryotic cells use an ensemble of covalent histone modifications to mark important sites of regulatory activity. One of these marks, trimethylation of lysine 36 on histone 3 (H3K36me3) is an abundant and highly conserved chromatin modification that is enriched at gene bodies of transcriptionally active genes and also at centromeric regions^1–3^. The presence and distribution of H3K36me3 domains at actively transcribed genes is conserved from human to yeast, suggesting that this mark is extremely important for proper cellular function^4^. Important regulatory activities linked to the H3K36me3 mark include transcription elongation^5,6^, prevention of cryptic start sites^7^, as well as pre-mRNA splicing^8^ and processing^9^. In addition to transcription, H3K36me3 also plays important roles in the recruitment of DNA repair machinery to mismatch regions, and for this reason, actively transcribed genes pre-marked with H3K36me3 are especially protected from DNA damage^10^.

The genome-wide distribution of H3K36me3 is maintained through various mechanisms. In human cells, H3K36 is mono- and di-methylated by eight distinct histone methyltransferases; however, the predominant writer of the trimethyl mark on H3K36 is SETD2^1,11,12^. Interestingly, SETD2 is a major tumor suppressor in clear cell renal cell carcinoma^13^, breast cancer^14^, bladder cancer^15^, and acute lymphoblastic leukemias^16–18^. In these settings, mutations and deletions of SETD2 often leads to global downregulation of H3K36me3 and are strongly associated with decreased overall survival^13–19^. H3K36me3 is enzymatically removed from chromatin by members of Jumonji domain-containing histone demethylase protein families^20^. However, the mark may be also removed by other mechanisms, including histone turnover. Histone turnover is performed by diverse ATP-dependent chromatin remodeling complexes, such as BAF (SWI/SNF) and related complexes^21,22^, as well as histone chaperones (e.g. FACT complex^23^ or SPT6^24^), that catalyze histone eviction from chromatin. Interestingly, histone turnover is highest at active promoters and less pronounced within transcribed regions enriched for H3K36me3^25,26^. In addition to the writers and erasers of this mark, H3K36me3 is also read by a range of the proteins harboring the PWWP domain as well as Tudor, chromo and plant homeodomain (PHD) finger domains^27^, many of which are known transcription regulators or DNA damage responders (e.g. PHF1^28^, Dnmt3a^29^ and LEDGF/p75^30,31^).

Transcription-coupled H3K36me3 deposition at coding regions is facilitated through the unstructured yet highly conserved C-terminal domain of RNA polymerase 2 (RNAP2 CTD). During transcription elongation, the RNAP2 CTD recruits many factors to transcribed genes, including the histone methyltransferases MLL (KMT2A; H3K4 methylation), DOT1L (H3K79me2), and SETD2 (H3K36me3) (**Figure 1A**)^32–34^. These methyltransferases are recruited to sites of transcription in part via direct interaction with RNAP2 ^6,35–38^. However, this common mode of recruitment curiously does not result in similar patterns of histone modification, as each of these marks has a distinct, characteristic profile across the genome^39–42^. For the H3K36me3 mark, serine-2 phosphorylation on the CTD of the POLR2A (Rpb1) subunit of RNAP2 interacts with the transcription elongation factor and histone chaperone SPT6, which associates with SETD2 through its binding partner IWS1^6,43,44^. Through this association (**Figure 1B**), H3K36me3 is thought to be propagated along gene bodies in a transcription-dependent manner^35^. In yeast, transcription is sufficient for formation of H3K36me3 domain over gene bodies^45^. Here the presence of SPT6 and IWS1 is not only essential for SETD2 recruitment, but also for the catalysis of H3K36me3 placement^6^. However, in mammals, SETD2 associates with a large number of nuclear proteins and complexes that possess their own chromatin recognition domains (e.g. p53^46^ or STAT1^47^), which could potentially influence SETD2 methyltransferase activity independently of transcription. Furthermore, it has been proposed that the placement of H3K36me3 involves “spreading” of the mark across the gene body^48–50^, and a variety of differing patterns reported for H3K36me3 profiles across gene bodies^51–54^. It has therefore remained uncertain whether SETD2 can be recruited to chromatin to place and spread the H3K36me3 mark through transcription-independent mechanisms in addition to transcription-dependent interaction with the RNAP2 CTD. Given the importance of H3K36me3, we were surprised to find no systematic genome-wide studies describing the spatial distribution, intensity, and dynamics of this mark placed by SETD2 in concert with other factors.

**Figure 1.**
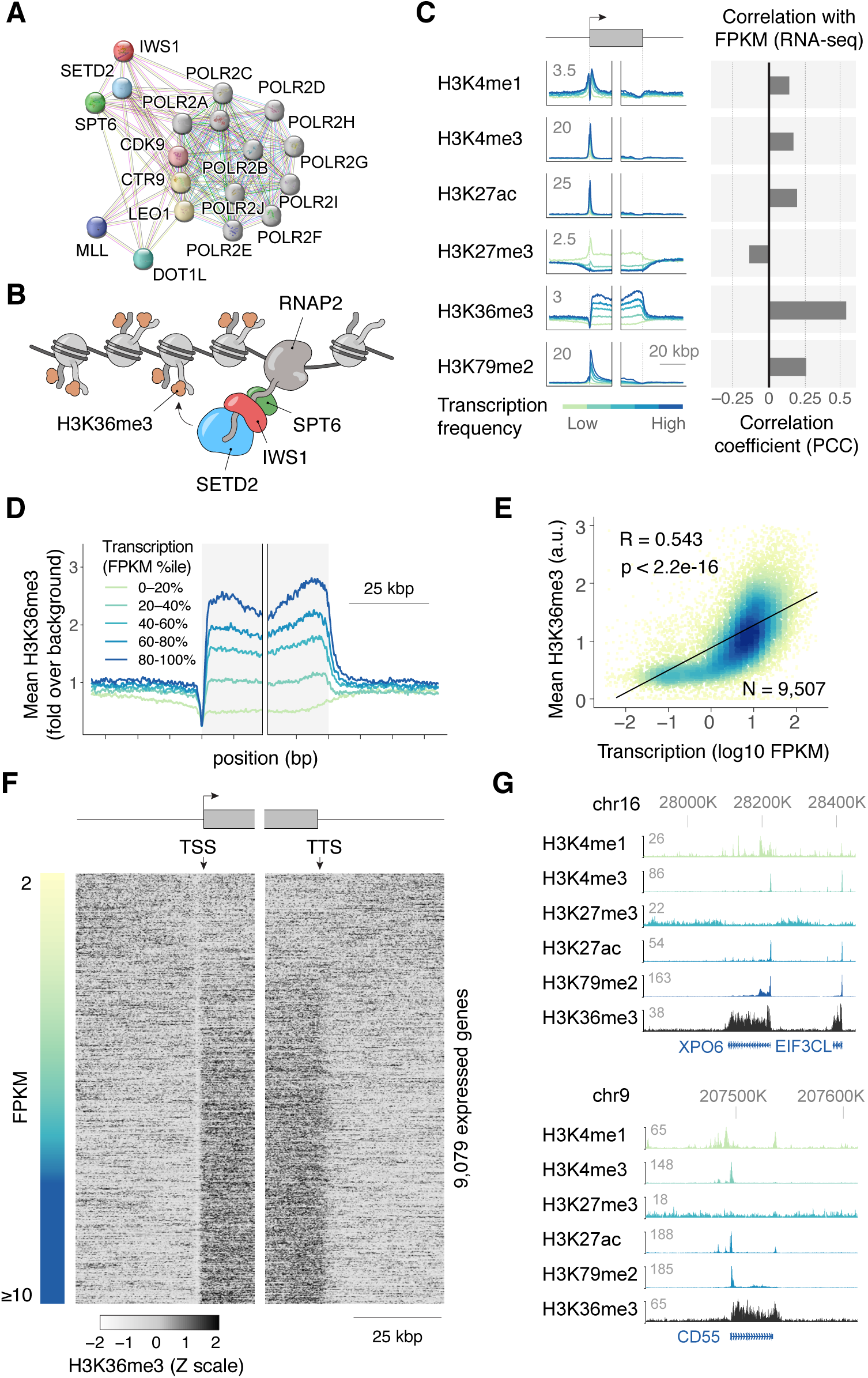
H3K36me3 placement by the SETD2:IWS1:SPT6 complex and genome-wide dependence on transcription. **(A)** STRING analysis of validated interactions between RNA Polymerase 2 (RNAP2) and histone methyltransferases MLL (KMT2A), DOT1L, and SETD2. The SETD2:IWS1:SPT6 ternary complex has extensive interactions with RNAP2 subunits. **(B)** IWS1 and SPT6 interact with the RNAP2 unstructured C-terminal domain to recruit SETD2, leading to H3K36me3 placement. **(C)** Analysis of histone ChIP-seq signals at gene bodies in K562 cells, organized by the degree of transcription measured via RNA-seq. H3K36me3 is the histone mark with the highest correlation to transcription. Measurement of correlation values is provided in **Figure S1. (D)** H3K36me3 profiles are flat plateaus over transcriptional units starting with the transcription start site (TSS) through the transcription termination site (TTS) in K562 cells. **(E)** The mean ChIP-seq enrichment of H3K36me3 scales with the transcript abundance measured with RNA-seq. **(F)** Heatmap of H3K36me3 plateaus at individual transcription units with their associated RNA-seq output measured by fragments per kilobase million (FPKM). **(G)** Individual genome browser tracks in K562 cells.

Although great effort has been made to elucidate the participation of H3K36me3 in various cellular processes^1^, much less is known about dynamics of the H3K36me3 mark itself in human cells. How are the activities of enzymes that regulate H3K36me3 writing and erasure coordinated to result in formation of these domains? Is the transcription-coupled H3K36me3 placement sufficient to achieve the steady-state of these domains, or are other transcription-independent processes needed to propagate and stabilize these marks? What are the insulation mechanisms that prevent this mark from spreading outside of its boundaries?

Because chromatin and its marks are highly dissipative complex systems^55,56^, non-linear and agent-based kinetic models have become invaluable tools to understand the dynamical behavior of the chromatin landscape and to predict the behavior of epigenetic systems^57–62^. We previously developed a model of H3K9me3 dynamics premised on simple nucleation-propagation-turnover Markov processes that provided quantitative predictions of kinetic properties and dynamics of H3K9me3 domains throughout the genome^58^. This model allowed us to elucidate the constraints that lead to the characteristic pattern of enrichment and memory of H3K9me3 domains in embryonic stem cells and fibroblasts^59^. Here we have further developed this approach to model the establishment and dynamic properties of H3K36me3 domains at gene bodies throughout the genome. Our analysis shows that H3K36me3 domains are broad, flat domains with sharp borders at gene bodies despite the lack of discernable insulator features in multiple cell types. We find that the abundance of H3K36me3 at gene bodies scales in proportion to transcription frequency, consistent with transcription-dependent marking by SETD2. In contrast to the direct local spreading of H3K9me3 marks in heterochromatin, our analysis shows that transcription-independent H3K36me3 spreading must be small in proportion to transcription-dependent mechanisms. Furthermore, incorporating measured rates of histone turnover is sufficient to produce remarkably in-vivo-like mark distributions over gene bodies without insulator elements. Finally, we find that SPT6 loss sharply reduces the observed dependence of H3K36me3 levels on transcription frequency, consistent with our model of H3K36me3 placement occurring primarily through transcription-dependent marking. Together our data provide new insights into the placement, persistence, and function of H3K36me3 domains.

## Results

### The quantitative dependence of H3K36me3 marking on transcription frequency

We began by performing a comprehensive analysis of histone mark ChIP-seq signals at gene bodies based on the degree of transcription. We analyzed the ChIP-seq profiles of H3K4me1, H3K4me3, H3K27ac, H3K27me3, H3K36me3, and H3K79me2 at gene bodies in K562 chronic myeloid leukemia cells. Despite extensive studies of the relationship between transcription and histone modification domains, we found few systematic reports relating the quantitative accumulation of all such marks to the frequency of transcription. We therefore sought to assess the relationship between the levels of each of these marks with transcriptional frequency, using the fragments per kilobase million (FPKM) levels from RNA-seq in the same cells as a proxy. We found that all of these marks highly statistically significant (p<2.2e-16) correlation with transcription. All marks positively correlated with transcription except for H3K27me3, which displayed a negative correlation (**Figure 1C and S1**). Of those with positive correlation, all histone marks except for H3K36me3 displayed a strong characteristic peak localized at transcription start sites (TSSs), consistent with regulatory roles during transcription initiation. In contrast, we found that H3K36me3 domains are localized directly over transcriptional units, with sharp, well-defined borders near TSSs and transcription termination sites (TTSs, **Figure 1D**), consistent with many previous studies of H3K36me3^9,19,40^. By comparing the mean levels of H3K36me3 across gene bodies to their FPKM values, we found that H3K36me3 had the highest correlation (R=0.543, p<2.2e-16) between transcriptional activity and histone mark levels over several powers of 10 (**Figure 1E**). Altogether, the unusually high correlation and presence of the mark throughout the entire transcriptional unit indicated a strong link between H3K36me3 accumulation and transcription elongation.

We next sought to address whether the shape of the H3K36me3 domain varied with transcriptional activity by analyzing these marks at individual loci genome-wide. We restricted our analysis to transcripts with FPKM values >2, and with gene bodies that spanned the 15-kb window of our analysis. The resulting heat map of 9,079 genes (**Figure 1F**) revealed that although the intensity of H3K36me3 marking varies with transcriptional activity, the overall shape of the distribution is largely unaffected, yielding flat plateaus at gene bodies throughout the genome. This interpretation was confirmed by examination of browser tracks of individual genes (**Figure 1G**). Nearly identical genome-wide results were obtained in A549 cells, an unrelated non-small cell lung cancer line, indicating these features are shared across multiple cell types (**Figure S2**).

### A transcription-dependent model of H3K36me3 dynamics

We next constructed a dynamic Markov state machine model to describe the dynamics of H3K36me3 marking and its relationship to transcription frequency. We constructed a one-dimensional lattice model of chromatin, analogous to the “beads on a string” picture of chromatin. Although higher-order chromatin structure may contribute to certain features of transcription, our relatively simple treatment of chromatin structure is justified, since well-ordered, evenly spaced arrays of nucleosomes are clearly visible at lowly and highly expressed genes in our analysis of MNase-seq data in K562 cells (**Figure S3**). In our simulations, the lattice {*I*_j_} spans from j = 1 to j = 512, where each lattice position *j* corresponds to an individual nucleosome. Thus with a nucleosomal spacing of 195 bp, our lattice model corresponds to 99.8 kbp of DNA. To make our model as general as possible, we consider that each nucleosome position along the lattice may occupy one of two states: an unmodified (*I*_j_ = 0) state or a modified with H3K36me3 (*I*_j_ = 1) state. Briefly, four features define the standard model (**Figure 2**):

**Figure 2.**
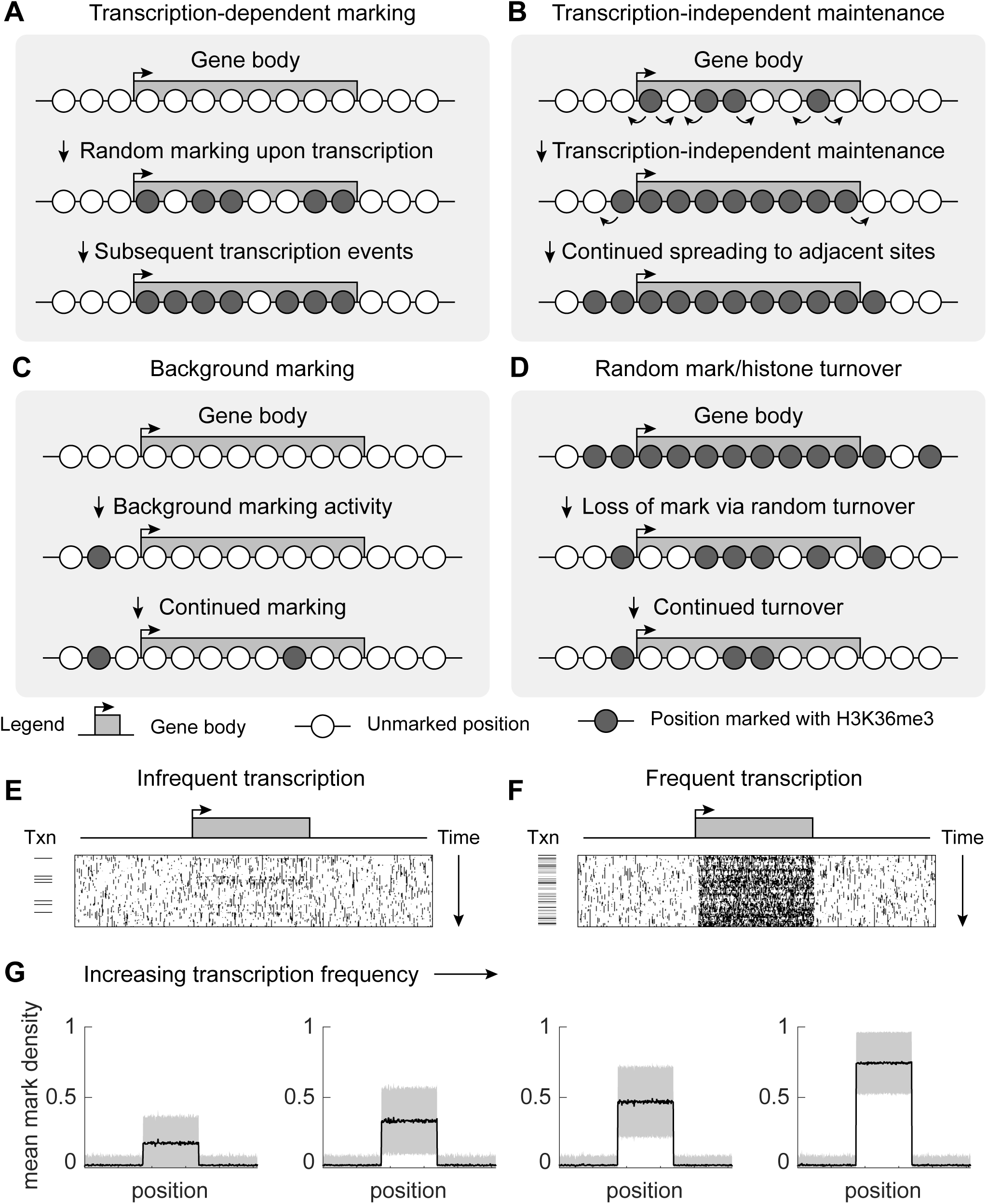
General components of the kinetic model of H3K36me3 at transcriptional units. **(A)** During a transcription event, nucleosomes within a transcriptional unit (gene body) are marked with constant probability. **(B)** H3K36me3 marks may be maintained by transcription-independent processes that lead to spreading of the marks to neighboring sites with constant probability. **(C)** All sites are marked with a very low background activity with constant probability. **(D)** Sites marked with HK36me3 are randomly turned over to unmarked sites with constant probability. **(E)** Visualization of an individual simulation over time reveals transient accumulation of H3K36me3 marks that are dissipated when transcription is infrequent. **(F)** Visualization of an individual simulation over time reveals steady-state levels of H3K36me3 marks are maintained when transcription is frequent. **(G)** The profile of simulated H3K36me3 levels in simulations scales in intensity with transcription frequency.

1. *Transcription-dependent marking*: Upon an individual transcription event occurring in an individual time step, nucleosome positions in a transcriptional unit (positions 174-338) are each randomly marked with probability *P*_n_. This process corresponds to the recruitment and stochastic labeling of nucleosomes during transcription events. Transcription events occur within a given time step at probability *P*_i_.
2. *Transcription-independent spreading*: Our general kinetic scheme integrates any processes that locally spread H3K36me3 into a single net propagation probability (*P*_+_). In our model, *P*_+_ describes the transcription-independent spreading of H3K36me3 to nucleosome positions immediately adjacent to H3K36me3-marked sites at each time step. This scenario represents the hypothetical pathway whereby SETD2 is recruited to sites of H3K36me3, for example directly or indirectly through H3K36me3 reader(s). Mechanistically, this process is likely composed of multiple rate-limiting steps (e.g. recruitment, marking), however for simplicity we treat this spreading as occurring as a single rate-limiting step.
3. *Background marking*: All nucleosomes across the lattice are subject to weak, low probability of marking at each time step. This probability is *P*_bg_, and corresponds to any process, such as non-specific SETD2 recruitment, that lead to non-specific marking.
4. *Random mark/histone turnover*: At each time step, all positions across the lattice are subject to stochastic removal of the H3K36me3 mark. All nucleosome marks are removed at each time step with probability *P*_–_. This process corresponds to any mechanism that results in the loss of the mark, and includes both H3K36 demethylation and histone/nucleosome turnover.

The implementation of this reaction scheme is performed using Monte Carlo simulation, where each of these processes occurs as a homogeneous Poisson process with exponentially distributed event times. The lattice space of the simulation is implemented with periodic boundary conditions, hence no insulator elements are present. Full details of this simulation are described in the *Methods* section, and MATLAB code of its implementation is provided as supplemental information.

When these processes are permitted to occur simultaneously, a dynamic H3K36me3 domain is established over the simulated gene body (**Figure 2E**). These simulated domains bear several features in common with our observations of H3K36me3 ChIP-seq data. In particular, simulations with low transcription probabilities bear elevated levels of H3K36me3 compared to background. Moreover, H3K36me3 levels of lowly transcribed genes in these simulations are significantly reduced compared to simulations with high transcription frequencies (**Figure 2E-G**).

To assess the contribution of the transcription-independent spreading, we performed 1,014 simulations by spanning a range of *P*_n_, *P*_i_, and *P*_+_, while keeping *P*_bg_ and *P*_–_ constant. The output of each of these simulations spanned a range of outcomes from very little marking throughout the simulation, to full saturation of the simulation space. Between these extremes, a domain of H3K36me3 is established that shows a strong dependency on transcription frequency. The full set of these outcomes is presented in **Figure S4**.

By analyzing the influence of the outcomes of the spreading parameter *P*_+_, we found that elevated mark spreading in a transcriptionally independent manner would result in poor establishment of H3K36me3 domains with sufficient contrast above non-transcribed regions (**Figure 3**). We found that the major effect of increasing the spreading term P+ was to increase background marking intensity without providing a corresponding increase in marking of the gene body. This outcome was true regardless of whether transcriptional frequency was low or high (**Figure 3A**).

**Figure 3.**
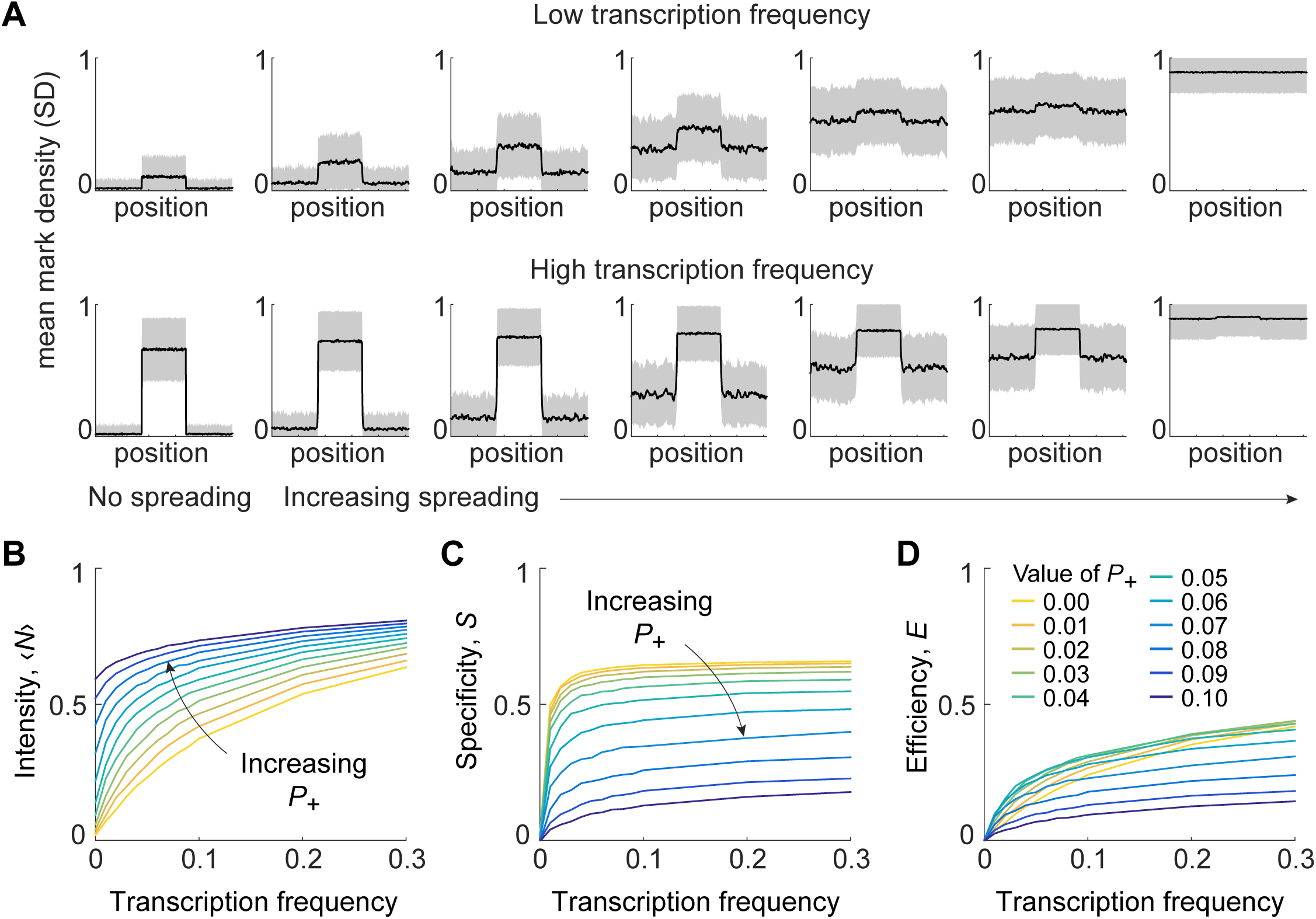
The effects of transcription-independent spreading of H3K36me3 in simulations. **(A)** For both low- and high-transcription simulations, as local spreading probability increases, the entire simulation experiences increased levels of marking, reducing the contrast of the mark between gene body and background. **(B)** Plot of mean H3K36me3 mark intensity at gene bodies as a function of transcription probability and spreading probability. **(C)** Plot of H3K36me3 specificity at gene bodies as a function of transcription probability and spreading probability. **(D)** Plot of H3K36me3 efficiency at gene bodies as a function of transcription probability and spreading probability.

To analyze this phenomenon in quantitative terms, we adopted a similar set of metrics to those we used previously to characterize H3K9me3 domains^58^. To quantify the mean intensity of marking, we use the term ⟨*N*⟩, which reports the fraction of nucleosome positions in the gene body with H3K36me3 marks. To quantify the overall specificity of the domain, we used a specificity score *S*:

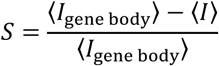

where ⟨*I*_gene body_⟩ is the mean mark density at the gene body, and ⟨*I*⟩ is the mean mark density over the entire lattice. Finally, the contrast between gene bodies and background regions is reflected by the efficiency score *E*:

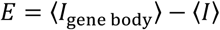

By plotting the intensity, specificity, and efficiency of the domains across different parameters, it is readily apparent why elevated spreading results in poor domains. While the intensity of marking increases within gene bodies, the specificity of this marking strongly drops due to the presence of additional marks outside of gene bodies (**Figure 3B-C**). The net result of elevated spreading is to reduce the contrast of the marks between the gene bodies and background regions (**Figure 3D**). Based on this result, we conclude that transcription-dependent is the dominant mode of H3K36me3 placement, and therefore anticipate that non-transcriptional propagation of H3K36me3 in cells is likely to produce a small fraction of the overall H3K36me3 marks at gene bodies.

### Replication-independent histone turnover induces an asymmetric distribution of H3K36me3 at the TSS

One notable feature of the H3K36me3 ChIP-seq tracks that was not captured by our model is the reduction of mark intensity near the TSS. This reduction is found in many individual tracks and is largely independent of transcription frequency (**Figure 1D**). We hypothesized that increased histone turnover in this region could induce a substantial reduction of H3K36me3 intensity in a dynamic manner by decreasing the effective lifetime of the H3K36me3 mark near the TSS.

We therefore sought a metric for histone turnover across the gene body. Several experimental approaches have been developed to measure the absolute kinetics of histone lifetime, including metabolic labeling and CATCH-IT ^63,64^. These methods have revealed a striking correspondence of histone lifetime with the profiles revealed by ChIP of the replication-independent histone H3.3^64^. We developed a variant of our standard model, where the rate of histone turnover was not uniform throughout the gene body, but varied by position (**Figure 4A**). We used the average profile of H3.3^65,66^ as a proxy for the relative rates of replication-independent histone turnover across the gene body and compared the results of the standard model (constant turnover) with the variable turnover model.

**Figure 4.**
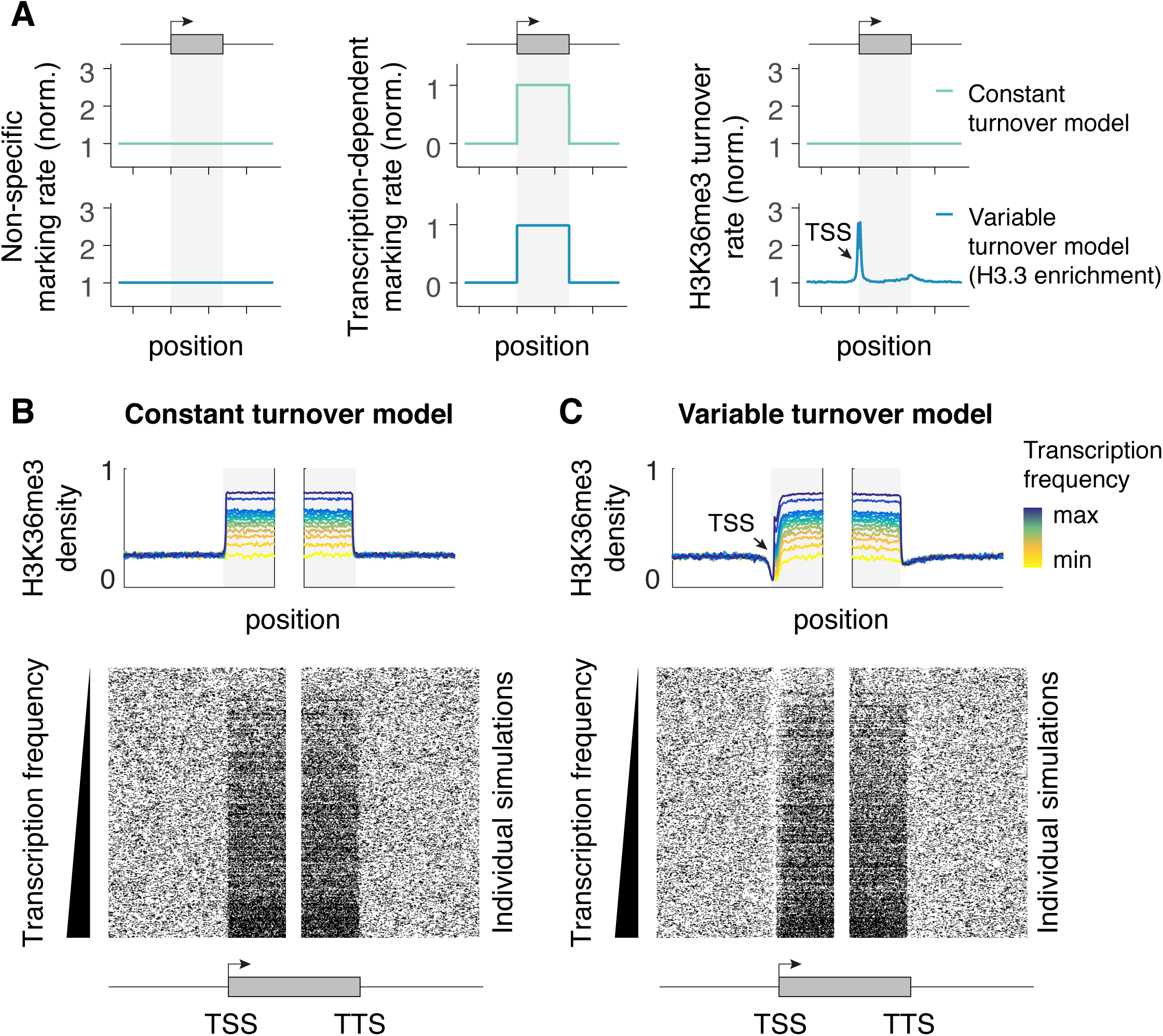
Variable histone turnover reproduces in vivo-like distributions of H3K36me3. **(A)** In addition to the standard constant turnover model, a variable turnover model where histone turnover is incorporated is presented. The rate of replication-independent histone turnover is obtained by measuring the profiles of H3.3 over gene bodies from ChIP-seq. **(B)** The constant turnover model reproduces essential features of H3K36me3 as observed from ChIP-seq data, including the observed scaling with transcription frequency. **(C)** The variable turnover model reproduces several in vivo-like features of H3K36me3 as observed from ChIP-seq data, including the characteristic asymmetric depletion and scalloped edges at the TSS, as well as the observed scaling with transcription frequency.

Compared to the standard model, the resulting H3K36me3 profiles from the variable turnover model are strikingly similar to the genomic profiles observed from ChIP-seq (**Figure 4B-C**). These profiles also show a similar dependence on transcription frequency as the ChIP-seq data. We therefore conclude that a full description of H3K36me3 dynamics requires including the varied rates of histone turnover that occur near transcription start sites. Our findings furthermore suggest that H3K36me3 is not faithfully copied nor rapidly spread onto the replication-independent H3.3 histones, as these processes would otherwise mask this dependency. Consequently our work confirms that the levels and genome-wide profiles of H3K36me3 are regulated in a replication-independent manner.

### SPT6 loss impairs the scaling relationship between H3K36me3 and transcriptional frequency

To validate our model and test whether transcription-coupled activities are responsible for most H3K36me3 placement, we sought experimental confirmation that disruption to the SETD2:IWS1:SPT6 ternary complex would affect the scaling relationship between H3K36me3 abundance and transcription frequency. We therefore compared mononucleosomal H3K36me3 signals obtained following siRNA transfection against either luciferase negative control (siLuc) or SPT6 (siSPT6), previously obtained elsewhere^67^. To measure transcription frequency, we used 3’ RNA-seq data from the same cells. Counts obtained using 3’ RNA-seq directly report on the frequency of transcription events at a given locus.

By plotting the H3K36me3 profiles based on transcription frequency in the siLuc control, we observed the scaling relationship between transcriptional frequency and H3K36me3 levels as described in Figure 1 above, confirming that the control does not affect this relationship (**Figure 5A-B**, R=0.462). In contrast, transfection of cells with siSPT6 causes a profound but incomplete reduction in the levels of H3K36me3 at gene bodies (**Figure 5C**). Moreover, this treatment alters the relationship between transcription frequency and H3K36me3 levels: Instead of observing a strong scaling relationship, the H3K36me3 levels show a reduced dependence on transcription frequency (**Figure 5C-D**, R=0.05). Pooling of transcripts into quintiles reveals that this effect persists despite pooling and across genomic replicates (**Figure 5E**) [F(1,14) = 13.4, p = 2.6e-3]. Together, our analysis shows that disruption of the SETD2:IWS1:SPT6 ternary complex alters the relationship between H3K36me3 accumulation and transcription frequency, providing an important validation of our kinetic model.

**Figure 5.**
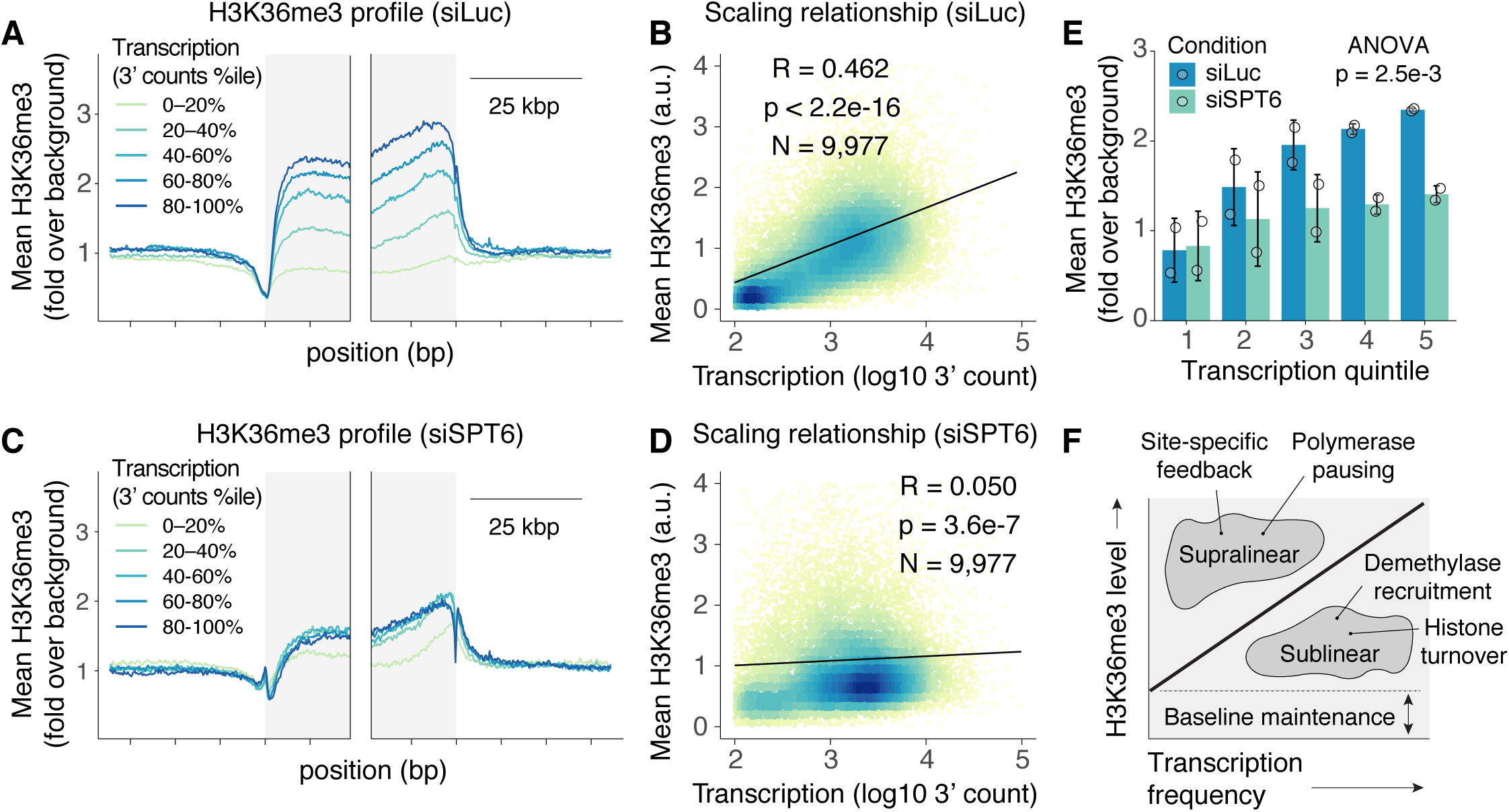
Validation of transcription-dependent kinetic model of H3K36me3 marking. **(A)** Analysis of the profile of H3K36me3 at gene bodies as a function of transcription frequency following control RNAi silencing of luciferase (siLuc). **(B)** Mean H3K36me3 level scales in the siLuc control with transcription frequency based on analysis of 3’ RNA-seq counts. **(C)** Analysis of the profile of H3K36me3 at gene bodies as a function of transcription frequency following RNAi silencing of SPT6 (siSPT6). **(D)** Mean H3K36me3 level scales in the siSPT6 with transcription frequency based on analysis of 3’ RNA-seq counts. **(E)** Analysis of transcription dependent H3K36me3 marking following segmenting transcripts into quintiles. Analysis from n=2 independent genome-wide replicates reveals a reduced scaling relation between transcription frequency and H3K36me3 levels. **(F)** Conceptual model describing the scaling relationship between transcription frequency and H3K36me3 levels. Deviations from this relation can arise from several sources of site-specific biological regulation.

## Discussion

Here we have employed quantitative approaches to accurately model the properties of H3K36me3 domains. Our comprehensive analysis of histone marks indicates that H3K36me3 domains are uniquely located throughout the transcribed regions of genes in different cell types. In contrast to some reports suggesting that H3K36me3 have strongly biased distributions over the gene bodies, our analysis shows that H3K36me3 domains are characterized as being broad, flat plateaus that span the entirety of gene bodies, with sharply defined boundaries and only modest deviations in intensity from the TSS to the TTS. We find a direct correspondence between transcription frequency and the intensity of H3K36me3 marking, indicating a tight relationship between transcription and H3K36me3 placement. Of the histone modifications we analyzed, the plateau-like distribution is characteristic only of H3K36me3, unlike other marks which show a high degree of enrichment near the TSS. The even marking throughout the gene body suggests that placement of the H3K36me3 mark occurs continuously throughout the entire process of transcription elongation.

Many questions have remained regarding the mechanisms that explain the accumulation of these marks across gene bodies. We used lattice-based simulation approaches to assess the contribution of transcription-dependent marking, transcription-independent maintenance (spreading) and histone turnover to the H3K36me3 domain shape. Our results indicate that transcription-independent spreading at even low rates would increase the H3K36me3 background levels outside of the coding regions. We find that elevated rates of spreading would eventually lead to collapse of the sharp domain boundaries that are observed in vivo. Therefore, our model suggests that transcription-dependent marking is the major pathway for H3K36me3 domain formation. This is in agreement with observation that H3K36me3 levels are proportional to the transcription frequency of individual genes. Importantly, SETD2-dependent H3K36me3 marking has essential roles at sites of double-stranded DNA breaks^68^. Although we find that spreading mechanisms do not contribute substantially to normal sites of transcription, we cannot currently exclude the involvement of transcription-independent spreading mechanisms during double-strand break repair.

Incorporation of replication-independent histone turnover into our model reduces the mark intensity near the TSS, yielding a strikingly similar domain shape to profiles measured in cells. Interestingly, genome-wide studies show that many active genes have a nucleosome-free region at the transcription start site. Our analysis (**Figure S5**) demonstrate that unlike H3K36me3 intensity, the size of this region does not correlate with transcription frequency, suggesting that the steady-state exclusion of histones from promoter regions is independent of H3K36me3 accumulation.

Transcription-dependent H3K36me3 marking relies on RNAP2:SETD2:IWS1:SPT6 complex formation^6^. Because our analysis suggested that co-transcriptional marking was the major pathway of H3K36me3 placement, we sought to test and validate this model by quantifying the effects following disruption of the SETD2:IWS1:SPT6 ternary complex. We analyzed ChIP-seq data obtained in the presence and absence of SPT6. Despite the loss of transcription scaling in SPT6 knock-down cells, a modest accumulation of H3K36me3 is preserved over gene bodies, suggesting that some kind of non-transcriptional mechanisms may still be operating in the absence of SPT6. As such, it is tempting to speculate that residual marking might reflect transcription-independent processes that may contribute to bookmarking of gene bodies independently of complexes involving SPT6. Such processes might be essential to prevent the complete silencing of genes by Polycomb complexes, since H3K36me3-containing nucleosomes are poor PRC2 substrates and H3K36me3 antagonizes PRC2-mediated H3K27 methylation^69,70^. In general, we propose that significant deviations of the transcription dependence of H3K36me3 levels may reflect biologically important sources of regulation, for example sites where polymerases move unusually quickly or slowly through a gene^71^, or where other regulatory factors such as H3K36 demethylases are in close proximity (**Figure 5F**)

Genome-wide analysis of H3K36me3 shows that it does not spread outside of the gene body boundaries, despite the lack of any obvious insulator elements. Furthermore, others have shown using an inducible system that these marks also do not spread onto adjacent loci^45^. In the same study, the authors found that the H3K36me3 mark persisted in the locus for 60 minutes after transcription inhibition, suggesting a short epigenetic memory for recently occurred transcriptional activity. This finding is consistent with our model, which suggests a short persistence of H3K36me3 in the absence of transcription. We therefore conclude that the strong dependence on co-transcriptional processes limits the capacity of the H3K36me3 mark to serve as a stable, heritable epigenetic mark at gene bodies.

Many of the processes dependent on H3K36me3 are ubiquitous and highly conserved from yeast to humans. Additionally, H3K36 oncohistone mutations^72^ and loss of the H3K36me3 methyltransferase SETD2^73^ both play important roles during cancer development. In particular, lysine-to-methionine mutations of H3K36 (H3K36M) contribute to distinct subtypes of sarcomas and pediatric chondroblastoma^74,75^. In these patients, expression of H3K36M mutants leads to global reduction of H3K36 methylation, including H3K36me3 throughout gene bodies, and shapes the oncogenic transcriptome of these cancer cells. Additionally, SETD2 deletions and loss-of-function mutations have been identified in patients with clear cell renal cell carcinoma (ccRCC), and alterations of SETD2 have been uncovered in pediatric high-grade gliomas, colorectal cancer and several hematologic malignancies. In ccRCC, progressive deregulation of H3K36me3 is linked to metastasis and is associated with overall lower survival ^19,76^. The H3K36me3 demethylases (JMJD2A and B) have also been implicated in colorectal cancer^77^ and ER positive breast cancer^78^ development. Overall, dysregulation of H3K36me3 homeostasis represents a common pathway that contributes to several malignancies.

Drugs targeting chromatin regulators are currently viewed as promising treatment avenues for various diseases, and the number of such drugs in clinical use is anticipated to grow in the foreseeable future^79–83^. Unfortunately, the cross-talk between chromatin modifying enzymes and other factors responsible for the maintenance of their modifications remains enigmatic. Insight into this interplay is an essential first step towards rational targeting of the chromatin landscape. Additionally, understanding of marking kinetics is invaluable for drug dosage and sustainability of the desired effect. Therefore, investigation of dynamic mechanisms and kinetic properties that underlie steady-state activities of each mark is crucial for the development of future treatments.

The chromatin landscape is a highly dissipative complex system that constantly integrates hundreds of signals and coordinates appropriate responses. Simulation approaches premised on simple assumptions like those that we present here enable modeling of different scenarios and are therefore ideal for quantitative profiling of the dynamical behavior and kinetics of chromatin marking. Additionally, a number of modern chemical biology tools and chemical probes enable rapid perturbation and characterization of highly dynamic and complex systems like the chromatin landscape for experimental validation of these processes^83^. Future models incorporating additional processes, such as co-transcriptional regulation, may further illuminate the functions of H3K36me3 and its role in development and disease.

## Acknowledgments

This work was supported by NIH grant R00CA187565 (H.C.H.), the Cancer Prevention & Research Institute of Texas Grant RR170036 (H.C.H.), the V Foundation grant V2018-003 (H.C.H.), and Gabrielle’s Angel Foundation for Cancer Research (H.C.H.), as well as GACR grants 16-06357S and 19-14360S (V.V.)

## Author contributions

K.C., V.V., and H.C.H. conceived of work, K.C. and H.C.H. analyzed data and wrote the manuscript. E.A.S., V.V., and H.C.H. provided critical feedback.

## Methods

### Processing of ChIP-seq data

Data set files for H3K4me1, H3K4me3, H3K27me3, H3K27ac, H3K79me2, and H3K36me3 were obtained as bigWig files from the ENCODE Consortium for both K562 and A549 cells. These files were provided by the ENCODE Consortium based on uniquely mapped reads to the hg19 genome build. H3.3 ChIP seq profiles from HeLa cells were obtained from GEO accession GSE45023. All data was processed and analyzed from at least 2 independent biological replicates.

### Heat maps and metagene plots

ChIP-seq bigWig files were processed using bwtool^84^ to extract the regions 34,600 bp upstream of TSSs, and 17,300 bp downstream of TSSs, and the resulting values were averaged at 200 bp windows. The resulting matrix was used to generate metagene profiles and heatmaps using R. Similar matrices were obtained using the same approach for the region 17,300 bp upstream of TTSs and 34,600 bp downstream of TTSs. Scaled whole-gene body metagene plots were not used because they were found to introduce biases that did not exist in unscaled plots.

### RNA-seq analysis and integration with ChIP-seq data sets

FPKM values were obtained directly from the ENCODE Consortium following mapping to the hg19 genome. To harmonize the relationship between RNA-seq and ChIP-seq data, the UCSC table browser was used to map annotations reported using Ensembl transcript identifiers to gene symbols. For RNA-seq, the longest transcript was chosen as the canonical transcript, and its TSS was used for ChIP-seq analysis. All analyses were performed by matching the gene symbols between ChIP-seq and RNA-seq data sets. All data was processed and analyzed from at least 2 independent biological replicates.

### MNase analysis and measurement of nucleosomal spacing

MNase-seq data was obtained from the ENCODE Consortium for K562 cells as bigWig files. For each canonical TSS defined above, we began by Z scaling the MNase-seq signal in the window ± 4 kbp from the TSS. The +1 nucleosome was then identified by the position where the Z-scaled MNase-seq signal reached 0.5 downstream of the TSS, and heat maps were created beginning 200 bp upstream of this position. Phase scores were assigned for each locus, and the top-scoring TSSs containing strong phases were used to plot the matrix of MNase signals. The resulting signal was averaged across all TSSs with sufficiently high phase scores and showed a readily apparent 195-bp periodicity. The precise value of this periodicity was measured with greater precision by examining the strongest frequency component following fast Fourier transform (FFT). The highest frequency signal in the FFT corresponded to 1/195 bp, confirming the canonical 195-bp periodicity reported elsewhere^85^.

### Protein-protein interaction network graphing

Protein-protein graphs of functional interactions between proteins were plotted using STRING^86^.

### Browser tracks

Genome browser tracks were generated using the WashU Epigenome Browser, with the resulting tracks exported as SVG files.

### Monte Carlo simulations

Implementation of Monte Carlo simulations was performed using custom code written in MATLAB (Mathworks, Natick, MA). All Monte Carlo steps were implemented using classic Monte Carlo methods and will be provided as supplemental data upon acceptance of the manuscript. Briefly, The stochastic trajectory of a given genetic locus is carried out by iteration of the following processes at each time step (see Figure 2):

1. Transcription-coupled marking: if transcription is occurring in a given time step, each gene body site at position j that is unmarked (I_j_ = 0) is converted to a marked state (I_j_ = 1) with probability P_n_ (where P_n_ = k_n_Δt). Transcription occurs within each given time step randomly with probability P_i_.
2. Transcription-independent spreading: for each marked nucleosome j on the lattice, if the j − 1 position is unmarked (I_j−1_ = 0), it is converted to a marked state (I_j−1_ = 1) with probability P_+_. Similarly, if the j + 1 position is unmarked (I_j+1_ = 0), it is converted to a marked state (I_j +1_ = 1) with probability P_+_. Mark propagation thus proceeds outward in both directions from each marked site.
3. Background marking: each position j that is unmarked (I_j_ = 0) is converted to a marked state (I_j_ = 1) with probability P_bg_ (where P_bg_ = k_bg_Δt).
4. Turnover: for each marked nucleosome j on the lattice, conversion to an unmarked state (I_j_ = 0) occurs with probability P_−_ (where P_−_ = k_−_Δt).
5. Time evolution: Simulation time t is incremented by Δt.

The above procedure treats transcription, marking, and turnover as homogeneous Poisson processes with exponentially distributed lifetimes. We also allow all simulations to evolve using periodic boundary conditions; therefore, there is no boundary implicit in the simulations. For all simulations, the value of Δt was chosen to be sufficiently small such that *P*_–_ = 0.05. Source code for Monte Carlo simulations will be provided before or upon acceptance.

### Statistical analyses

All statistical procedures were performed using R. All reported P values were obtained using two-sided tests. When calculated p-values are smaller than the 64-bit double precision machine epsilon (2^−52^ = 2.2e-16), p-values are reported as p<2.2e-16.

## Supplemental Figure Legends

**Figure S1: Relationship between transcription frequency and ChIP-seq enrichment of chromatin marks at gene bodies.** The mean ChIP-seq enrichment of **(A)** H3K4me1, **(B)** H3K4me3, **(C)** H3K27ac, **(D)** H3K27me3, **(E)** H3K36me3 and **(F)** H3K72me2 marks in K562 cells is plotted against the transcript abundance measured by RNA-seq.

**Figure S2: Analysis of histone marks at gene bodies in A549 cells. (A)** H3K36me3 profiles over transcriptionally active genes; transcription start sites (TSS) and transcription termination sites (TTS) are highlighted. The individual genes clustered by transcription abundance determined by RNA-seq. **(B)** The mean ChIP-seq enrichment of H3K36me3 scales with transcript abundance measured by RNA-seq. **(C)** Heatmap of H3K36me3 plateaus at individual transcription units with their associated RNA-seq output measured by fragments per kilobase million (FPKM). **(G)** Individual genome browser tracks for different chromatin marks in A549 cells.

**Figure S3: Analysis of nucleosome occupancy downstream of transcription start sites. (A)** Heatmap of MNase-seq fragments in K562 cells at individual genes aligned by their transcription start sites (TSSs) and arranged by transcription frequency. Arrays of nucleosomes positioned downstream of TSSs are highlighted. **(B)** Mean nucleosome density determined by MNase-seq aggregated across individual genes ordered by their transcription frequency. **(C)** Phasing of the MNase-seq signal is constant.

**Figure S4: Model output spanning all parameters.** Dependence of H3K36me3 domain shape and boundaries on frequency of transcription-independent spreading, background marking and transcription-dependent marking. The probabilities of transcription event (Pi), H3K36me3 propagation by transcription-independent spreading (P+) and by transcription-dependent placement (Pn) are underlying variables.

**Figure S5: Nucleosome-free regions do not scale with transcription frequency. (A)** Heatmap of MNase-seq fragments at individual genes aligned by their TSS and ordered by the size of nucleosome free region. The abundance of transcripts of individual genes measured by RNA-seq are highlighted. **(B)** The transcript abundance measured by RNA-seq plotted against size of nucleosome-depleted region (NDR). **(C)** H3K36me3 levels show no strong relationship with the size of the NDR.

