## Supplementary figures and images for "Dynamics of transcription-dependent H3K36me3 marking by the SETD2:IWS1:SPT6 ternary complex"

### Supplemental Figures

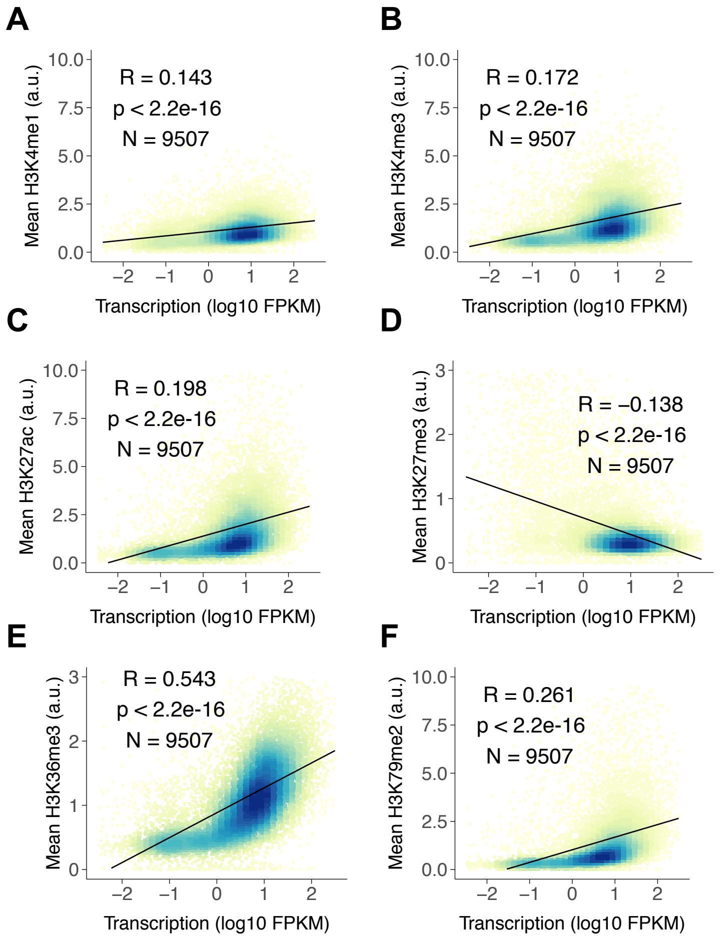

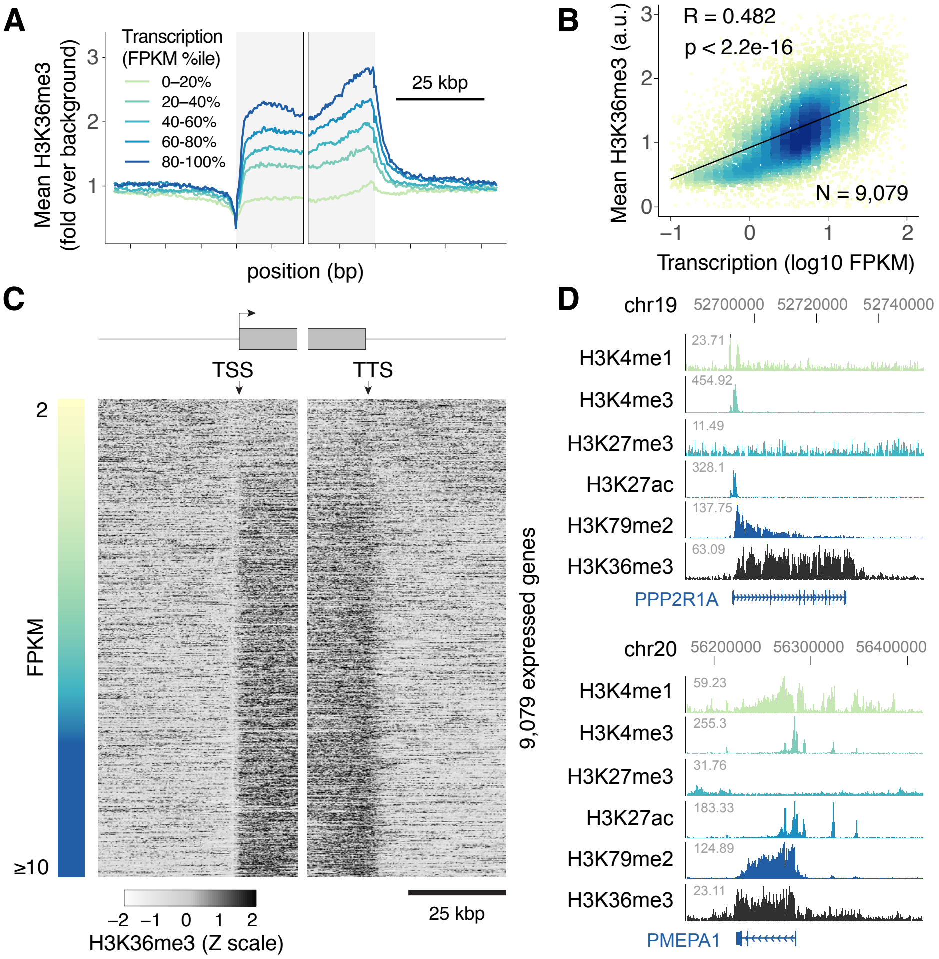

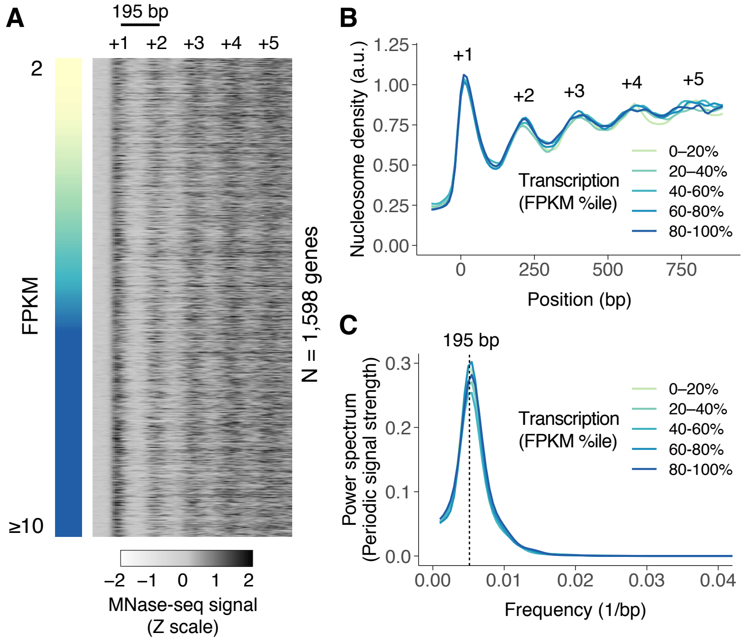

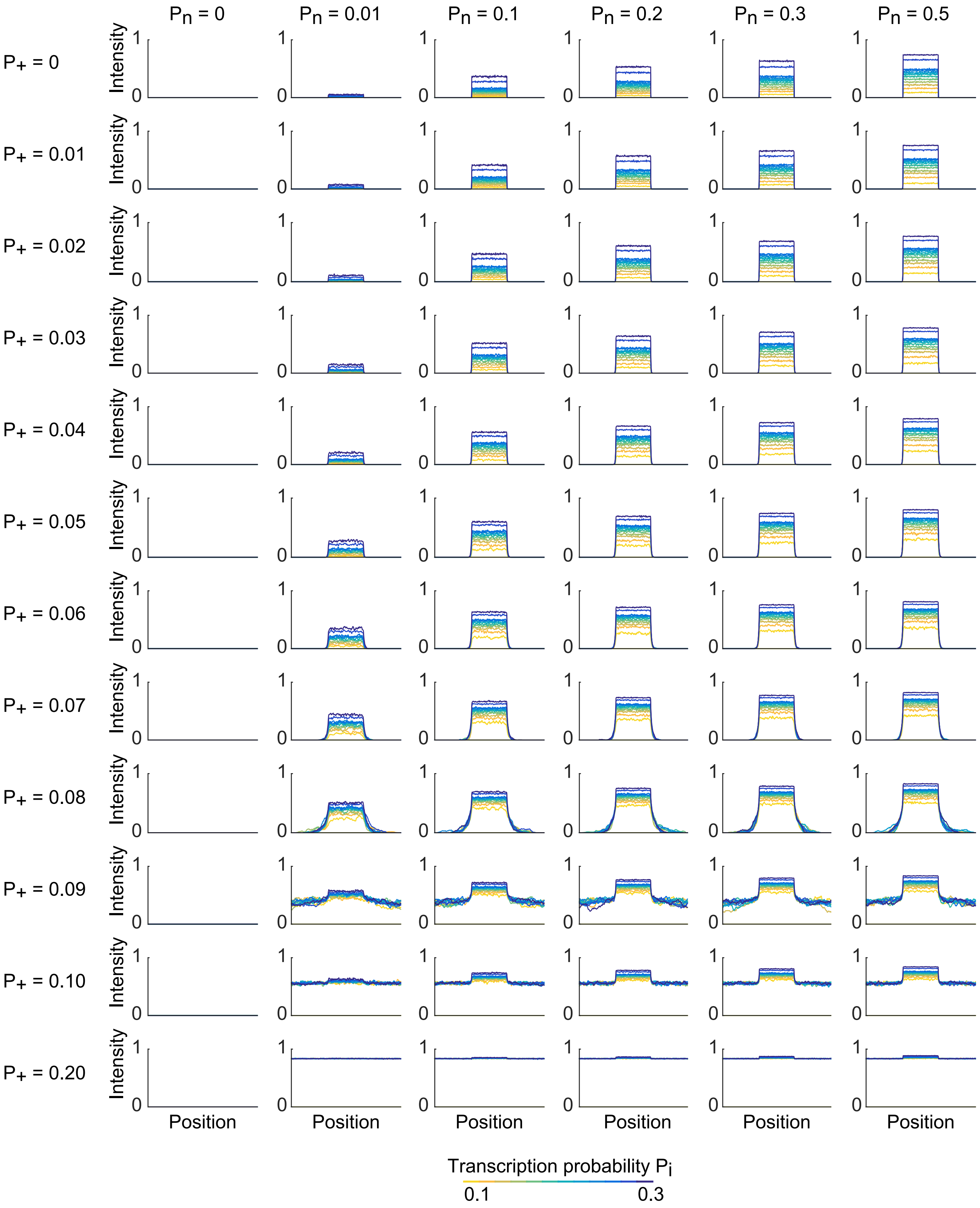

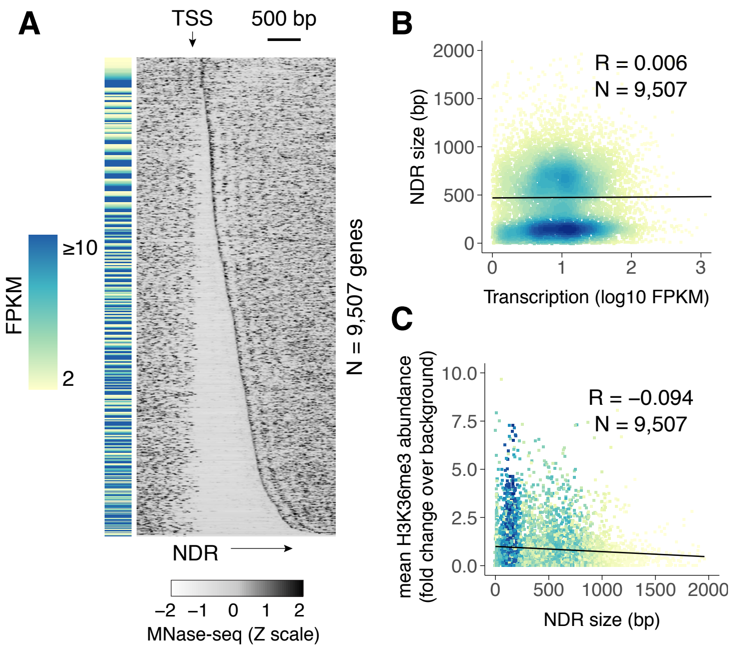
